# Comprehensive comparative analysis of the effects of temperature on the Notch signaling response *in vivo*

**DOI:** 10.1101/2025.04.18.649621

**Authors:** Nimmy S. John, Kah Seng Tang, Michelle A. Urman, ChangHwan Lee

## Abstract

Temperature is a critical factor that modulates cellular metabolism and stem cell regulation. Despite extensive studies, the influence of temperature on stem cell regulation via Notch signaling has been limited to studies relying on studies that involve indirect readouts to Notch activation. This study systematically analyzes the effects of temperature on the Notch signaling transcriptional response at the chromosomal, cellular, and tissue levels. Using complementary direct Notch readouts, we demonstrate that Notch activation remains largely unchanged across temperatures, suggesting the presence of temperature-compensatory mechanisms that maintain robust Notch activation. Notch transcriptional activity readouts, however, increased with temperature, indicating that elevated temperatures may enhance Notch transcriptional activity at the chromosomal level. These findings provide a comprehensive framework for understanding effects of temperature and offer new insights into the regulation of Notch signaling in stem cell biology.

## Introduction

Temperature is a crucial regulator of developmental timing and physiological homeostasis in both ectothermic and endothermic organisms^1^. Fluctuations in temperature can lead to short-term and long-term cellular changes, such as alterations in cell composition and proteins folding^2^. Studies have shown that temperature modulates cellular metabolism, influencing the rate of gene expression essential for organismal development^1,3,4^. It also plays a vital role in stem cell regulation by affecting signaling pathways that govern the viability, proliferation, and differentiation of stem cells^5^.

Notch Signaling is a conserved cell signaling pathway that operates via a common mechanism across metazoans, playing a pivotal role in regulating cell fate decisions and tissue patterning^6–10^. In *Drosophila*, Notch signaling remains stable across temperature fluctuations due to temperature-dependent compensatory mechanisms^11–13^. In contrast, in chick amniote brains, short-term hypothermia has been shown to enhance Notch activity and suppress neurogenesis in neural progenitor cells^1^. However, most of these studies rely on indirect readouts, such as GFP driven by Notch responsive promoters, highlighting the need for more direct and sensitive assays to assess how temperature modulates Notch activation and signaling dynamics.

Here, we focus on Notch activation in the *Caenorhabditis elegans* germline, where it directly drives transcription of two Notch targets, *sygl-1* and *lst-1*, to maintain a pool of 30-75 germline stem cells (GSCs) at the distal end of the gonad^14–16^. Temperature modulates multiple aspects of *C. elegans* physiology, including developmental speed, locomotion patterns, egg-laying rates, and chemosensory behaviors^17–21^. Notably, an approximate 5ºC increase in growth temperature accelerates developmental timing by about 50%, whereas lowering temperature is associated with extended lifespan^14,17,22^. Despite these broad effects, how temperature influences Notch activation and its functional consequences in the germline remains poorly understood. Here, we address this gap by using direct readouts of Notch activity to systematically analyze the effects of commonly used worm growth temperatures (15ºC, 20ºC, 22.5ºC, and 25ºC) on Notch-dependent transcriptional response.

Here, we used smFISH to perform a comprehensive comparative analysis of Notch-induced transcription across temperatures (15°C, 20°C, 22.5°C and 25°C). The spatial pattern and overall level of Notch signaling response remains largely unchanged across temperatures, suggesting the presence of temperature-compensatory mechanisms that maintain robust Notch activation. Individual ATS intensities and cytoplasmic mRNA counts, however, increased with temperature, indicating that elevated temperatures may enhance Notch transcriptional activity at the chromosomal level.

## Results

To systematically analyze the effects of temperature on Notch signaling response in the *C. elegans* germline, we performed single-molecule RNA fluorescence *in situ* hybridization (smFISH) to visualize transcripts of a major Notch target, *sygl-1*, in young-adult wild-type (N2) worms grown at different temperatures (15°C, 20°C, 22.5°C and 25°C) (Fig. 1A). *sygl-1* smFISH assays have been established as direct readouts of Notch-induced transcriptional activation and its spatial pattern^8,14,23–26^. These readouts include *sygl-1* nascent transcripts at active transcription sites (ATS), reflecting chromosomal-level Notch activity, and mature cytoplasmic mRNAs, estimating cellular-level activity ^8,14,23,25–28^. Notch activity at 20°C was consistent with previously reported measurements across chromosomal, cellular, and tissue levels^14,23,24,26^ (Fig. 1, 20ºC).

**Figure 1:**
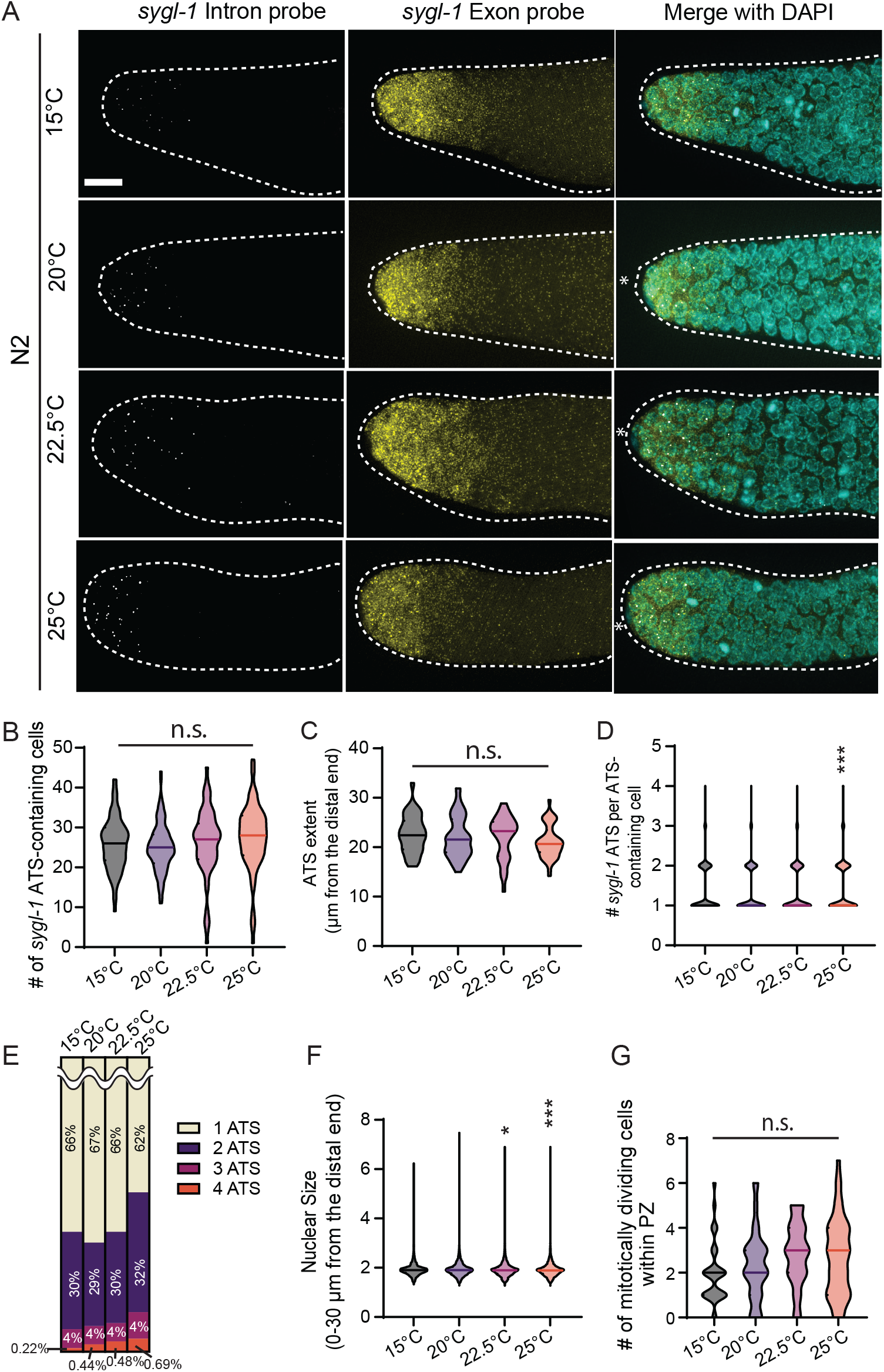
Overall changes in *sygl-1* transcriptional response as temperature increases. (A) Z projected *sygl-1* smFISH images that are representative of the *sygl-1* Notch response as temperature increased (15°C, 20°C, 22.5°C and 25°C). (B) # of *sygl-1* ATS-containing cells were plotted as a population for each temperature. (C) *sygl-1* ATS Extents were plotted as a population for each temperature. Sample sizes for temperatures, 15°C, 20°C, 22.5°C and 25°C were as follows: n =35,56, 36, and 47 gonads respectively. (D) The number of *sygl-1* ATS per ATS-containing cells was plotted as a population for each temperature. (E) Percentages of 1-4 *sygl-1* ATS distribution were plotted as a population for each temperature. (B, D-F) Sample sizes for temperatures, 15°C, 20°C, 22.5°C and 25°C were as follows: n =86,118,63, and 84 gonads respectively. (F) Nuclear sizes via radius with 30μm were plotted as a population for each temperature. Sample sizes for temperatures, 15°C, 20°C, 22.5°C and 25°C were as follows: n =6091,10330,4899 and 6782 nuclei respectively. (G) Mitotic cell counts were plotted as a population for each temperature. Sample sizes for temperatures, 15°C, 20°C, 22.5°C and 25°C were as follows: n = 35, 56, 36, and 45 gonads respectively.

### Notch-induced transcriptional activation remains unchanged across different temperatures

To compare tissue-level Notch-induced transcriptional activation across temperatures, we scored the number of cells containing *sygl-1* ATS and the extent of *sygl-1* ATS along the gonadal axis, which reflects the overall Notch transcriptional response and the size of Notch-responsive germ cell pool, respectively (Fig. 1B-C). Both measurements remained consistent across all temperatures, indicating that Notch activation is unaffected by temperature at the tissue level (Fig. 1B-C). To assess Notch activation at the cellular level, we analyzed the number of *sygl-1* ATS in each ATS-containing cell (Fig. 1D-E, S1A-B). Neither the ATS number nor the composition of ATS per cell varied with temperatures, further confirming that Notch activation remains robust across temperatures at both cellular and tissue levels (Fig. 1D-E). Supporting this, the distributions of germ cell nuclear sizes, indicative of cell cycle stage, were essentially the same across temperatures, suggesting normal cell cycle progression regardless of temperature (Fig. 1F, S1C). However, we observed an increase in mitotically dividing cells at higher temperatures (Fig. 1G), consistent with previous reports of accelerated cell cycle progression at elevated temperatures, such as 25ºC^3,29^. Altogether, these results demonstrate that Notch-induced transcriptional activation remains unchanged at cellular and tissue levels across physiological temperature ranges.

### Transcriptional activation of a Notch-independent gene, *let-858*, increases with temperature

We next asked whether the temperature-independent consistency in transcriptional activation is a unique feature of Notch target genes, or a broader property shared with Notch-independent genes. To address this, we performed smFISH targeting *let-858* (Fig. 2A), a gene whose expression is independent of Notch signaling in germ cells^14,23,24,30,31^. We focused our analysis on the first 30 μm of the distal gonad, where the majority of GSCs reside^14,32^ and quantified the number of *let-858* ATS (Fig. 2B). In contrast to the *sygl-1* ATS, the number of *let-858* ATS increased with temperature (Fig. 1B-D and 2B). These results suggest that temperature-independent transcriptional consistency is specific to Notch-induced transcriptions and does not extend to all actively transcribed genes.

**Figure 2:**
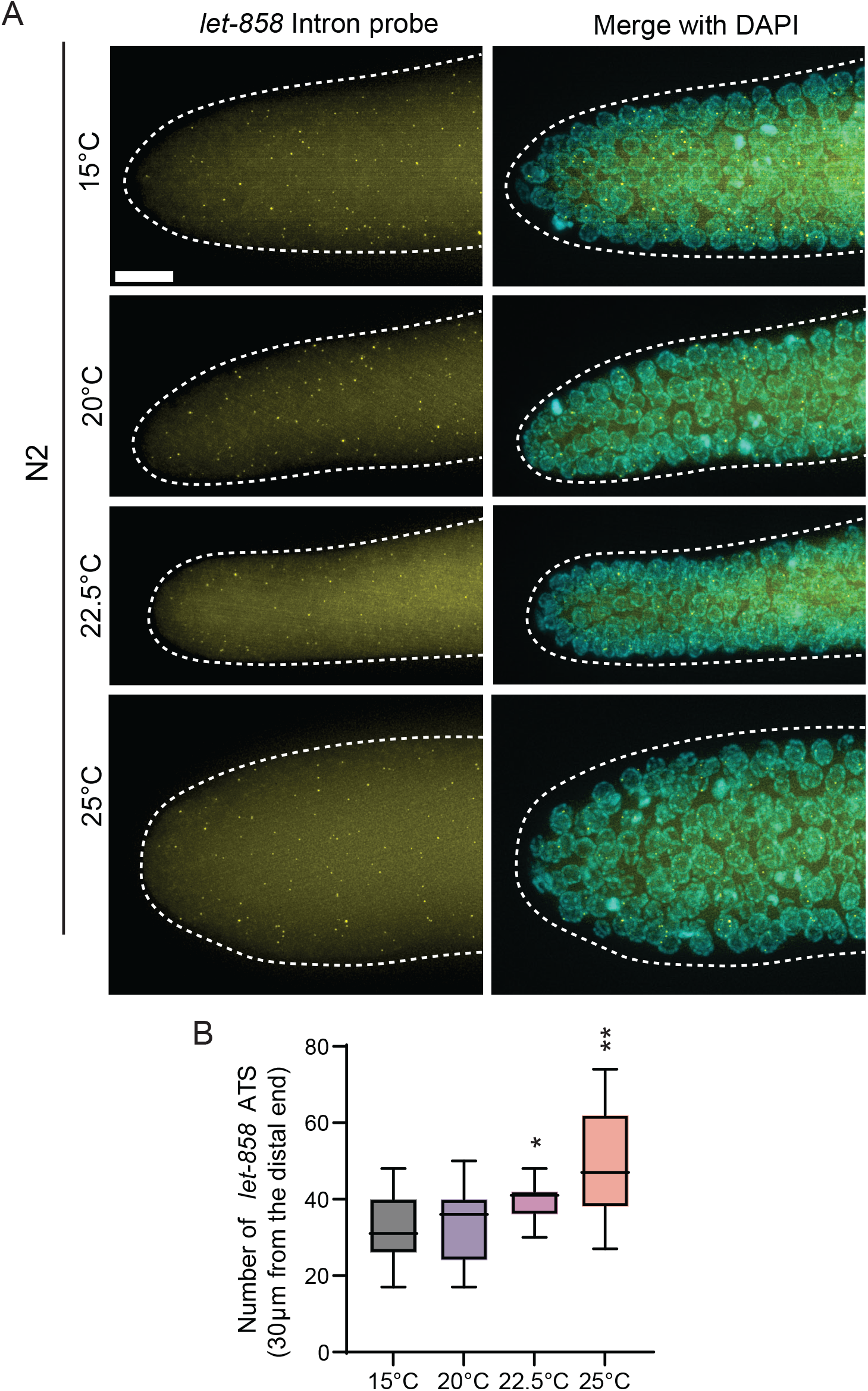
Effects of temperature on the *let-858* transcriptional response. (A) Z projected representative *let-858* smFISH images as temperature increases. (B) Number of *let-858* ATS within 30 μm from the distal end was plotted as a population for each temperature. Sample sizes for all temperatures were n = 15.

### Notch-induced *sygl-1* transcriptional activity increases with temperature

Although Notch-induced transcriptional activation, which reflects the number of GSCs or chromosomes responding to Notch signaling, remains unchanged across temperatures (Fig. 1), we asked whether transcriptional activity, defined as the amount of RNA produced, is affected by temperature. To assess *sygl-1* transcriptional activity at the chromosomal level, we measured individual *sygl-1* ATS intensities at different temperatures and observed a gradual increase with rising temperature (Fig. 3A). Similarly, the summed *sygl-1* ATS intensity per nucleus, a proxy for cellular-level transcriptional activity, also increased with temperature (Fig. 3B). This trend extended to the tissue-level as both the total number of *sygl-1* mRNA within the first 60 μm of the distal gonad (approximately two-thirds of the progenitor zone, PZ) and the average number of *sygl-1* mRNAs per cell increased with temperature (Fig. 3C-D). The PZ size, an estimate of gametogenesis capacity, also expanded at higher temperatures, although the trend did not precisely mirror changes in ATS intensities or mRNA levels (Fig. 3E). Together, these results indicate that while Notch-induced transcriptional activation is buffered against temperature changes, Notch activity increases with temperature at both chromosomal and cellular levels.

**Figure 3:**
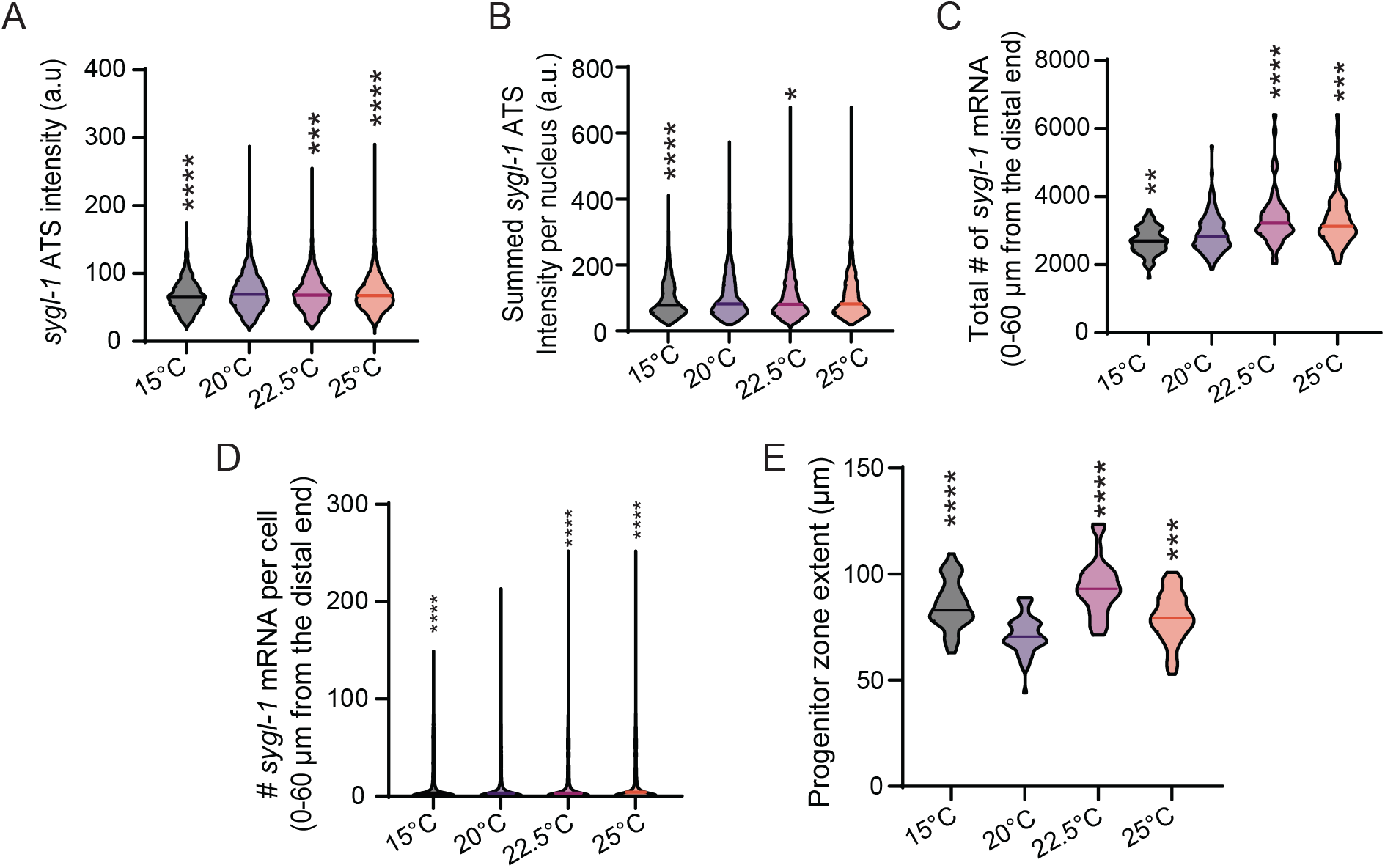
Overall changes in the Notch-dependent generation of cytoplasmic mRNA. (A) *sygl-1* ATS intensity was plotted as a population for each temperature. Sample sizes for temperatures, 15°C, 20°C, 22.5°C and 25°C were as follows: n =3256,4372,2467 and 3416 nuclei respectively. (B) Summed *sygl-1* ATS intensity was plotted as a population for each temperature. Sample sizes for temperatures, 15°C, 20°C, 22.5°C and 25°C were as follows: n =2223, 2954, 1654, and 2305 nuclei respectively. (C) Total number of *sygl-1* mRNA within 60μm from the distal end was plotted as a population for each temperature. Sample sizes for temperatures, 15°C, 20°C, 22.5°C and 25°C were as follows: n =86,118,63 and 84 gonads respectively. (D) Number of *sygl-1* mRNA per cell within 60μm from the distal end was plotted as a population for each temperature. Sample sizes for the temperatures, 15°C, 20°C, 22.5°C and 25°C were as follows: n = 15198, 27271, 12676 and 16896 nuclei respectively. (E) Progenitor zone extents were plotted as a population for each temperature. Sample sizes for temperatures, 15°C, 20°C, 22.5°C and 25°C were as follows: n = 35, 56, 36, and 45 gonads respectively.

### The spatial distribution of Notch-induced *sygl-1* transcription is unaffected by temperature changes

Notch-induced transcriptional activation occurs in a steep gradient within the GSC pool at the distal gonad, which plays a crucial role in germline polarization and GSC maintenance^14,23,24^. The spatial pattern of *sygl-1* ATS has also been established as a reliable indicator of the Notch-responsive GSC pool^14,23,24,32^ (Fig. 4A, red-dashed lines). To determine whether temperature influences this graded Notch response pattern and alters GSC pool size, we analyzed the spatial patterns of *sygl-1* ATS and mRNAs across temperature (Fig. 4). Specifically, we quantified the percentage of germ cells containing *sygl-1* ATS as a function of distance from the distal end of the gonad, which reflects the probability of Notch activation along the gonadal axis (Fig. 4A). Across all temperatures, the sygl-1 ATS gradient and the inferred GSC pool size remained largely unchanged, indicating that the spatial pattern of Notch activation is also buffered against temperature changes (Fig. 4A, red-dashed lines). This temperature-independent spatial pattern was also evident when analyzing the number of *sygl-1* ATS or number per cell and the percentage of germ cells with *sygl-1* mRNAs above the basal level (∼5 *sygl-1* mRNAs per cell) (Fig. 4B-D). These results indicate that the graded pattern of Notch-induced transcription and the size of the GSC pool are preserved across a range of physiological temperatures.

**Figure 4:**
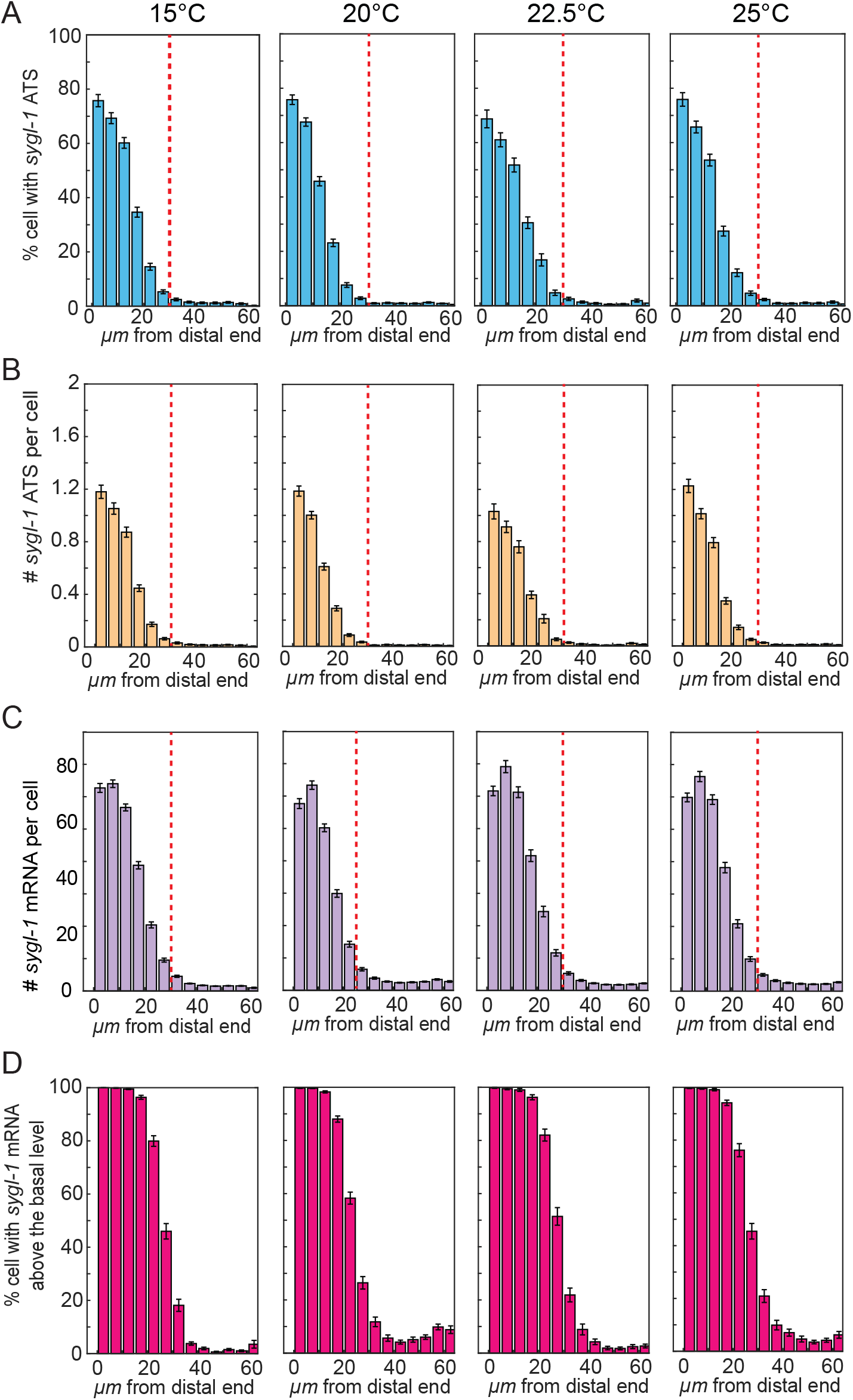
*sygl-1* transcriptional response is not affected spatially as temperature increases. (A) Percentages of cells with *sygl-1* ATS were plotted against the function of position from the distal end in microns for N2 in varying temperatures. (B) Number of *sygl-1* ATS per cell was plotted against the function of position from the distal end in microns for N2 in varying temperatures. (C) The number of *sygl-1* cytoplasmic mRNA per cell was plotted against the function of position from the distal end in microns for N2 in the varying temperatures. (D) Percentages of cells with *sygl-1* cytoplasmic mRNA above the basal level were plotted against the function of position from the distal end in microns for N2 in varying temperatures. (A-D) Sample sizes for temperatures, 15°C, 20°C, 22.5°C and 25°C were as follows: n =86, 118, 63, and 84 gonads respectively.

## Discussion

This study systematically investigates how physiological temperatures affect Notch-induced transcription at chromosomal, cellular, and tissue levels in the *C. elegans* germline. Using direct, quantitative Notch readouts, including *sygl-1* ATS and mRNAs, we show that the probability of Notch activation and its spatial pattern are remarkably invariant across a range of temperatures (15°C, 20°C, 22.5°C, and 25°C). In contrast, the transcription of the Notch-independent gene *let-858* increases with temperature (Fig. 2), consistent with previous reports showing temperature-sensitive regulation of transgene expression^33–35^. These findings suggest the presence of a buffering mechanism that maintains consistent Notch-induced transcriptional activation, likely to preserve GSC pool size and function under varying environmental conditions and ensure robust gametogenesis.

Despite stable Notch activation patterns, the transcriptional activity of *sygl-1*, as measured by individual ATS intensities and total mRNA per cell, increased with temperature (Fig. 3A-D). This enhanced Notch transcriptional activity correlates with expansion of the PZ (Fig. 3E), which may ultimately impact fertility and progeny size. We speculate that increased activity or abundance of transcriptional regulators, such as the DNA-binding protein LAG-1/CSL or the Notch intracellular domain (NICD), may underly this temperature-associated boost in *sygl-1* transcription. Further studies should assess whether temperature modulates the levels, binding dynamics, or nuclear accessibility of these key transcriptional components.

Notably, the temperature-buffering mechanism appears to regulate Notch transcriptional activation (i.e., the probability of Notch signaling response; Fig. 4A-B), but not the extent of transcriptional activity (i.e., the amount of RNA produced once activated; Fig. 4C-D). This distinction implies that buffering occurs upstream of target gene activation, potentially at the level of Notch ligand-receptor interactions. We speculate that temperature may influence the abundance or stability of Notch ligands (e.g., LAG-2/DSL) or modulate proteolytic processing of the GLP-1 receptor to release NICD. Elucidating how these molecular steps are insulated from temperature changes will be critical for understanding the robustness of Notch signaling in dynamic environments. Together, our findings reveal that while Notch activation is buffered against temperature changes, its transcriptional activity and output are more responsive, providing a layered regulatory architecture. This dual mode of control may allow for stable GSC regulation while enabling physiological flexibility in response to changing conditions.

## Materials and Methods

### Nematode strains used in this study

Nematode strains used in this study:
| Strain name | Genotype |
| --- | --- |
| N2 | Wildtype strain of *C. elegans* |

### Nematode culture

All strains were maintained at 20°C as previously described^36^. The wild-type was N2 Bristol. For the smFISH experiments, all strains were synchronized via hypochlorite treatment and cultured on OP50-seeded NGM plates until the appropriate day of adulthood within each temperature.

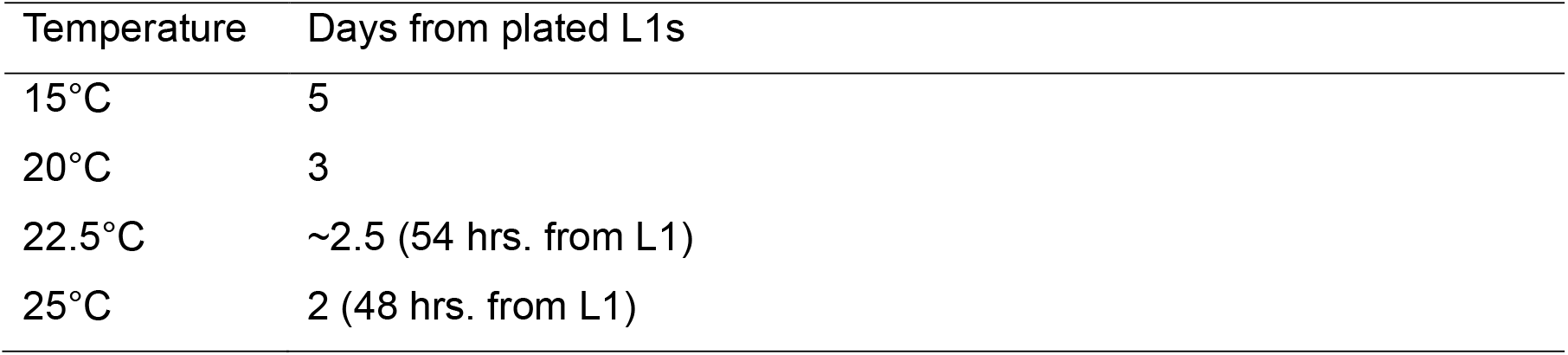

### Single-molecule RNA fluorescence *in situ* hybridization (smFISH)

smFISH for *sygl-1* and *let-858* were performed as previously described^14,23,24,37^. Synchronized L1 larvae were grown on OP50 until day 1 of adulthood within their respective temperatures as described in Nematode culture table above ^24,25^. Briefly, the synchronized *C. elegans* of N2 for each experimental set were washed off plates with 2-3 mL non-RNase free 1X PBS + 0.1% Tween-20 (PBST) and were collected on the 60 mm petri dish cover. An additional 2-3 mL of non-RNase free PBST was added, and the worms were dissected to extrude the gonads in PBST with 0.25 mM levamisole added. The dissected samples were fixed with 3.7% formaldehyde in 1X PBS with 0.1% Tween-20 at room temperature (RT) for 30 min, with rotation. Samples were spun down at 2000 RPM for 1 min unless noted otherwise. After fixation, samples were permeabilized with the permeabilization buffer for 10 min at room temperature with rotation. The samples were then washed twice with RNase free PBST, resuspended in 70% ethanol, and stored overnight at 4°C.

Custom Stellaris FISH probes (Biosearch Technologies, Inc., Petaluma, CA) were designed against the exon and intron regions of *sygl-1* and the intron regions of *let-858* as described previously^14,23^. Ethanol was removed and samples were incubated in 1 mL of wash buffer for 5 min at room temperature. Gonads were hybridized with 1 μL of each of the *sygl-1* probes (6.25 μM) or *let-858* probes (6.25 μM) in hybridization buffer for 24 hours at 37°C with rotation. After probe addition, samples were kept in the dark for all incubations and washes. Samples were rinsed once with wash buffer at room temperature, then incubated in wash buffer for 30 min at room temperature with rotation. The DNA was then labeled by incubation in smFISH wash buffer containing 1 mg/mL diamidinophenylindole (DAPI) for 30 min at room temperature followed by two short washes with smFISH wash Buffer. Finally, samples were resuspended in 10-12 μL Antifade Prolong Gold mounting medium (Life Technologies Corporation, Carlsbad, CA) and mounted on glass slides, which were then cured for 48 hours.

### Microscopy setup and image acquisition

Gonads were imaged completely (depth >15 μm) with a Z-step size of 0.3 μm using a Leica DMi8 Widefield Microscope that is equipped with a THUNDER Imaging system that computational clearing methods provided in the Leica Application Suite X (LAS X) acquisition software (Leica Microsystems Inc., Buffalo Grove, IL) as previously described^23,24^. All imaging was done with LED8 light sources, sequentially through the channels in decreasing wavelengths to avoid bleed-through and prevent any photobleaching from occurring. The illumination and exposure settings for the acquisition of the gonad images were set up as previously described^24^.

### Progenitor zone extents

In this study, the progenitor zone (PZ) extent was measured from the most distal end of the gonad to the end of the progenitor zone, where a cell row with more than one crescent-shaped cell as previously described^23,24,38–43^.

### Notch *sygl-1* ATS and mRNA extents

The *sygl-1* ATS extents and mRNA extents were measured as previously described to estimate the GSC pool size^23,24^. The ATS extents were measured using the distance from the distal most end of the germline to the last ATS within the germline. The mRNA extents were measured using the distance from the distal most end of the germline to the end of the mRNA-rich region (<5 mRNA per cell) within the germline^23,24^.

### Image processing using the custom-made MATLAB codes

All processes were implemented and automated using modified MATLAB (v2.0) codes similar to the source code developed in our previous work ^8,14,23,24,26^. After the analysis was completed, MATLAB and GraphPad Prism were used to visualize the data generated and conduct statistical tests as previously described^23,24^. If the datasets met the requirements for parametric statistical analysis through normality tests (Anderson-Darling Normality test), ANOVA and t-tests were performed to compare datasets presented in this study. If the data set did not satisfy the requirements for parametric analysis, the Kolmogorov-Smirnov (KS) test (a nonparametric version of the t-test) was performed.

## Acknowledgments

We are thankful for the resources provided by the Molecular Biology Core Facility in the Life Sciences Research Building and the RNA Institute at the University at Albany. This work was funded by the University at Albany (FRAP-A Award 1189585-1-97969).

## Figure Legends

**Figure S1:** (A) The number of *sygl-1* ATS per ATS containing cell within 30μm were plotted as a population for each temperature. (B) The number of *sygl-1* ATS per ATS containing cell within 20μm were plotted as a population for each temperature. (C) Nuclear Size via radius within 60μm were plotted as a population for each temperature. (D) Total number of *sygl-1* mRNA within 60μm from the distal end were plotted as a population for each temperature. (E) The number of *sygl-1* mRNA per cell within 60μm from the distal end were plotted as a population for each temperature

