## Supplementary figures and images for "Comprehensive comparative analysis of the effects of temperature on the Notch signaling response *in vivo*"

### Supplementary Figure 1

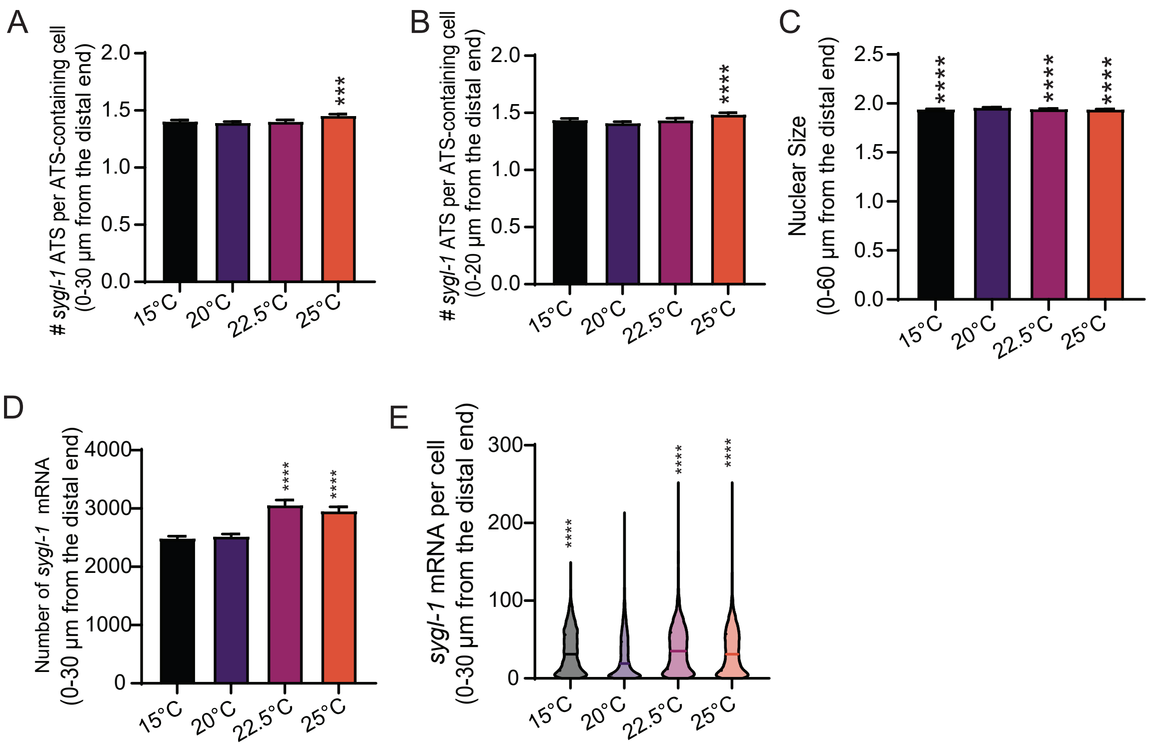
